# Small extracellular vesicular transfer of MYCN and glycolytic cargo coordinates metabolic and immunological reprogramming in neuroblastoma in vitro

**DOI:** 10.64898/2026.04.27.721043

**Authors:** Lin Ma, Mi Liu, Olga Piskareva

## Abstract

Extracellular vesicles (EVs) are emerging mediators of oncogenic communication within the neuroblastoma (NB) tumour microenvironment (TEM). Here, we investigated how constitutive MYCN overexpression influences the proteomic and functional properties of small EVs (sEVs) derived from SKNAS-MYCN-GFP (SK-M) cells and assessed their impact on non-cancerous immune cells. SK-M cells exhibited robust MYCN upregulation at both the mRNA and protein levels and produced sEVs that were selectively enriched in MYCN. Transwell co-culture revealed transfer of MYCN-GFP to recipient DC2.4 nuclei, indicating intercellular transport of functional transcription factor cargo. LC-MS/MS profiling showed that SK-M sEVs incorporated oncogenic cargo non-randomly, displaying significant enrichment of metabolic and MYC/MYCN-regulated pathways, including glycolysis, mTORC1 signalling, and suppression of oxidative phosphorylation (OXPHOS). These observations are consistent with emerging evidence that MYC family proteins can regulate metabolism through vesicular transfer of glycolytic kinases to neighbouring cells. Functionally, SK-M cells displayed elevated lactate secretion and reduced acetyl-CoA, and their sEVs induced a glycolytic shift in recipient immune cells, increasing lactate output in DC2.4, RAW264.7, BMDCs, and splenocytes. sEV-treated BMDCs and splenocytes acquired immunoregulatory phenotypes characterised by increased IL-10, reduced IL-12, expansion of regulatory T cells (Tregs), and macrophage polarization toward an M2-like state. These findings demonstrate that MYCN-driven NB cells disseminate metabolic and immunosuppressive cues via sEVs, reshaping the local immune landscape to favour tumour tolerance. This study provides mechanistic insight into how MYCN-amplified NB cells exploit EV-based communication to coordinate metabolic rewiring and immune escape.

## 1. Introduction

Neuroblastoma (NB) is the most common extracranial solid malignancy of childhood is characterised by profound clinical and biological heterogeneity(1–3). Among its molecular drivers, *MYCN* amplification remains the most powerful adverse prognostic factor, defining a high-risk subset with poor long-term survival despite intensive multimodal therapy(4–8).

Beyond its canonical roles in promoting proliferation, inhibiting differentiation, and sustaining self-renewal(5, 9, 10), MYCN profoundly remodels tumour cell physiology by orchestrating metabolic rewiring, oxidative and proteotoxic stress responses, and adaptive pro-survival signalling(11–14). MYCN amplifies aerobic glycolysis, alters mitochondrial dynamics, suppresses OXPHOS, and enhances anabolic pathways to meet the biosynthetic demands of rapidly dividing NB cells(11–14).

Recent evidence indicates that these metabolic programmes can extend beyond cell-intrinsic regulation. MYC family oncogenes, including MYCN, have been shown to regulate metabolism through vesicular transfer of glycolytic kinases, enabling cancer cell to disseminate metabolic traits to neighbouring cell via extracellular vesicles (EVs)(15). EVs, particularly small EVs (sEVs), are increasingly reccognised as key mediators of intercellular signalling within the NB tumour microenvironment (TME)(16, 17). They carry proteins, lipids, metabolites, mRNA, and miRNA that can reprogramme stromal components, enhance tumour invasion, and promote metastatic dissemination(18, 19). In NB, sEVs derived from tumour cells influence mesenchymal stromal cells, endothelial cells, and other components of the TME by delivering cytokines, miRNAs, and oncogenic mediators, thereby fostering a tumour-supportive niche(20–23).

However, the role of MYCN-enriched sEVs in shaping immune cell behaviour within NB TME remains poorly understood. Immune dysfunction, characterised by dendritic cell (DC) impairment, Treg expansion, and tumour-associated macrophage polarisaton, is a hallmark of high-risk NB and contributes to its classification as an “immune-cold” tumour(20, 24).

Metabolic shifts within the TME, particularly elevated lactate and acidic pH, further suppress cytotoxic immunity while promoting immunoregulatory phenotypes(25–27). Whether MYCN-driven sEVs participate in orchestrating this metabolic-immune axis has not been definitively explored(28).

In this study, we employed a constitutive MYCN-overexpressing NB model (SKNAS-MYCN-GFP, SK-M) to (i) investigate the incorporation and intercellular transfer of MYCN via sEVs, (ii) characterise MYCN-dependent proteomic alterations in sEVs, and (iii) determine how MYCN-driven sEV cargo remodels the metabolism and immune phenotype of recipient immune cells, including DCs and splenocytes. Our findings reveal that MYCN-overexpressing NB cells disseminate metabolic and immunosuppressive programmes via sEVs, promoting lactate-driven immunoregulation and fostering a tumour-tolerant immune niche. These insights provide a mechanistic link between MYCN oncogenic signalling, EV-mediated metabolic communication, and immune escape in NB.

## **2.** Materials and Methods

### 2.1 Cell lines

Human neuroblastoma SKNAS and murine macrophage RAW264.7 cells were maintained in Dulbecco’s Modified Eagle’s Medium (DMEM; Gibco#11966025, Grand Island, NY, USA) supplemented with 10% Fetal Bovine Serum (FBS; Gibco#10270106, Grand Island, NY, USA) and 1% Penicillin-Streptomycin (P/S; 5000U/mL; Gibco#15140122, Grand Island, NY, USA). The murine dendritic cell line DC2.4 was cultured in RPMI-1640(Gibco#21875034, Paisley, UK) supplement with 10% FBS and 1%P/S. All cell lines were obtained from the American Type Culture Collection (ATCC) and authenticated by DNA profiling before use.

### 2.2 Plasmid transfection

The MYCN-GFP plasmid (Public Protein/ Plasmid Library (PPL), China) encodes *MYCN* fused to enhanced GFP. 8 x 10^4^ SKNAS cells were transfected with Lipofectamine™ 2000 (Invitrogen^TM^, Cat#11668027, Carlsbad, CA, USA) using optimised plasmid-to-reagent ratios. Transfection efficiency was evaluated by fluorescence microscopy and confirmed by Western blotting for MYCN and GAPDH.

### 2.3 Generation of stable SKNAS-MYCN-GFP cells

Stable clones were generated by G418 (Procell, Cat#PB180125, Wuhan, China) selection. The optimal concentration was determined by titration (400 µg/mL achieved ∼80% cell death in untransfected SKNAS cells). Transfected cells were selected by 4 days, expanded, and cryopreserved.

### 2.4 Single-cell cloning

Post-selection, SKNAS cells were serially diluted and seeded at limiting density in 96-well plates. Wells containing a single GFP-positive cell were expanded, and MYCN expression was validated by Western blotting and quantitative PCR (qPCR).

### 2.5 Extraction of RNA and qPCR

Total RNA was isolated using TriQuick Reagent (Solarbio, Cat*#* R1100, Beijing, China). RNA concentration and quality were determined by NanoDrop™ 2000 (Thermo Scientific™, Cat#ND-2000, USA). Complimentary DNA (cDNA) was synthesised using a commercial reverse transcription kit (APExBIO, Cat#K1074, USA). qPCR was performed on a CFX96 Real-Time PCR system (Bio-Red) with MYCN and GAPDH primers (Supplementary Table S1).

Relative expression was calculated by the 2^-ΔΔCt method.

### 2.6 Cell culture for conditioned medium

SKNAS-GFP(SK) and SKNAS-MYCN-GFP(SK-M) cells (2.5 x 10^7^) were plated in T175 flasks with DMEM containing 10% exosome-depleted FBS (Gibco#A2720801, Grand Island, NY, USA) and 1% Antibiotic-Antimycotic (100X) (Gibco#15240062, Grand Island, NY, USA).

Conditioned medium was collected after 3 days of culture.

### 2.7 Proliferation and viability assay

Cell proliferation was assessed using the Quant-It™ PicoGreen™ dsDNA Assay Kit (Invitrogen, Cat#P7589, Eugene, OR, USA). Cell viability was determined using Cell Counting Kit-8 (CCK-8) (Sigma-Aldrich, Cat#96992, Waltham, Massachusetts, USA), and relative viability was expressed as fold change compared with control SK cells.

### 2.8 sEV isolation

EVs were isolated from conditioned medium by differential ultracentrifugation. Briefly, media were centrifuged at 16,000g for 30 minutes at 4⁰C to pellet microvesicles (mEVs), followed by ultracentrifugation at 110,000 x g for 90 min. The sEV pellet was washed in filtered PBS and underwent a second centrifugation at 110,000g before resuspension in PBS or RIPA buffer.

### 2.9 Nanoparticle tracking analysis (NTA)

sEV size distribution and concentration were determined using a Nanosight NS300 system (Malvern Instruments, Great Malvern, UK). Samples were diluted in filtered PBS to obtain an optimal concentration of 40-80 particles per frame. NTA was conducted at a temperature range of 20-25⁰C using a 488nm blue laser and a camera level set to 14. A sCMOS camera captured three-second videos of the sEVs in suspension, operating with a shutter speed of 1,000, gain of 400, and a frame rate of 25 fps. Data processing was performed using NTA software version 3.2 Dev Build 3.2.16, with a detection threshold set to 5, while parameter such as maximum jump mode, jump distance, and minimum track length were adjusted automatically.

### 2.10 Transmission electron microscopy (TEM)

For TEM, sEV were deposited onto 200-mesh copper grids coated with formvar and silicon monoxide (Ted Pella, Cat#01830, Redding, CA, USA), stained with uranyl acetate alternative (Ted Pella, Cat#19485, Redding, CA, USA), and imaged on a Hitachi H7650 transmission electron microscope (HITACHI, Tokyo, Japan) at 100 KV.

### 2.11 Confocal microscopy for MYCN transfer

To investigate intercellular *MYCN* transfer, DC2.4 cells (1.0 x 10^6^) were co-cultured with SK-M cells (5.0 x 10^4^) in a 0.4µm Polyethylene Terephthalate (PET) co-culture insert (Millipore, Cat# MCHT06H48, Molsheim, France). After 4 days, DC2.4 cells were fixed, stained with Hoechst dye (Beyotime, Cat#33342, Shanghai, China), and imaged on a Nikon Eclipse 90i microscope equipped with a Nikon Ri1 camera (Nikon, Tokyo, Japan).

### 2.12 Protein extraction and Western blotting (WB)

sEVs and cells were lysed in RIPA buffer supplemented with protease inhibitor cocktail (100X) (Thermo Scientific^TM^, Cat#78430, Rockford, IL, USA). Protein concentrations of sEV and cell lysates were determined by Micro BCA™ Protein Assay Kit (Thermo Scientific) (mBCA)(Thermo Scientific™, Cat#23235, Rockford, IL, USA) *a*nd Pierce BCA Protein Assay Kit (Thermo Scientific™, Cat#23225, Rockford, IL, USA), respectively, following the manufactures guidelines. Protein samples were diluted 4X Bolt™ LDS sample buffer (Invitrogen^TM^, Cat#B0008, Carlsbad, CA, USA) and supplemented with the appropriate volume of 10X Bolt™ Sample Reducing Agent (Invitrogen™, Cat#B0009, Carlsbad, CA, USA), and denatured at 95℃ for 5 min. Equal concentrations of protein (10 µg for cells, 5 µg of sEVs) were separated on 4-12% Bis-Tris plus gels (Invitrogen^TM^, Ca#NW04120, NW04122, NW04125, Carlsbad, CA, USA) and transferred onto 0.22-μm Polyvinylidene Difluoride membrane (PVDF) membranes (ThermoFisher Scientific™, Cat. # 88018, Rockford, IL, USA). Membranes were blocked in 5% bovine serum albumin (BSA) solution prepared in TBS-T (Tris buffered saline with 0.1% Tween-20) and incubated primary antibodies (Supplementary Table S2) overnight at 4℃, followed by HRP-conjugated secondary antibodies in TBS-T; anti-mouse IgG, HRP (Cell Signalling Technology, Cat#7076S, dilution1:3000, Danvers, MA, USA), or anti-rabbit IgG, HRP (Cell Signalling Technology, Cat#7074S, dilution1:3000, Danvers, MA, USA) for 60 min at room temperature. Each blot was developed using ECL Chemiluminescent Substrate Reagent Kit (Invitrogen^TM^, Cat#WP20005, Carlsbad, CA, USA) and imaged on the Amersham Imager 600 (Freiburg, Germany).

### 2.13 Mass Spectrometry

Proteins were extracted from cells and sEVs for quantitative and proteomic analysis. For digestion, 40 µg of total protein was mixed with five volumes of pre-chilled acetone and precipitated at −40 °C for 1 h. The pellets were collected by centrifugation at 12,000 rpm for 10 min at 4 °C, dissolved in 8 M urea, 50 mM Ammonium bicarbonate and sequentially treated with 5mM DTT, 20mM Iodoacetamide (IAA), and trypsin according to the manufacturer’s protocol, including a 2 h enzymatic digestion at 37℃. After digestion, peptides were desalted using spin columns and the final eluates (∼200 μL) were concentrated under vacuum and reconstituted in mobile phase A for LC–MS analysis.

For each sample, 200 ng of total peptides were separated and analyzed with a nano UPLC (nanoElute2) coupled to a timsTOF Pro2 instrument (Bruker) with a nano-electrospray ion source. Separation was performed using a reversed phase column (PePSep C18, 1.9 μm, 75 μm × 15 cm, Bruker, Germany). Mobile phases were H2O with 0.1% formic acid (FA) (phase A) and acetonitrile (ACN) with 0.1% FA (phase B). Separation of sample was executed with a 45 min gradient. The mass spectrometer adopts DDA PaSEF mode for DDA data acquisition, and the scanning range is from 100 to 1700 m/z for MS1. During PASEF MS/MS scanning, the impact energy increases linearly with ion mobility, from 20 eV (1/K0 = 0.6 Vs/cm2) to 59 eV (1/K0 = 1.6 Vs/cm2).

The mass spectrometry proteomics data have been deposited to the ProteomeXchange Consortium (https://www.iprox.cn/page/project.html?id=IPX0014697000) via the iProX partner repository with the dataset identifier PXD071984 (and iProX accession IPX0014697000).

### 2.14 Data processing

Vendor’s raw MS files were processed using SpectroMine software (4.2.230428.52329) and the built-in Pulsar search engine. MS spectra lists were searched against their species-level UniProt FASTA databases (uniprot_Homo sapiens_9606_reviewed_2024_05. fasta), Carbamidomethyl [C] as a fixed modification, Oxidation (M) and Acetyl (Protein N-term) as variable modifications. Trypsin was used as proteases. A maximum of 2 missed cleavage(s) was allowed. The false discovery rate (FDR) was set to 0.01 for both PSM and peptide levels. Peptide identification was performed with an initial precursor mass deviation of up to 20 ppm and a fragment mass deviation of 20 ppm. All the other parameters were reserved as default.

The resulting data were imported into Perseus software (version 2.0.3.1) for further processing. In Perseus, proteins identified by only a single peptide, by a single site, or those flagged as reverse hits or contaminants were excluded. LFQ values were log2(x) transformed and filtered for valid values. For protein identification, LFQ values were required to have at least three valid values for either the SK or SK-M samples. Differentially expressed proteins between SK and SK-M were identified after imputing, missing values from a normal distribution.

### 2.15 Ethics of Animal Experiments

All animal experiments were approved by the Animal Care and Use Committee of Soochow University (Approved ethics number:202404A0424) and conducted in strict accordance with ARRIVE guidelines (https://arriveguidelines.org).

6-week-old female C57BL/6 mice (SPF grade, body weight 18-20 g) were purchased from Shanghai SLAC Laboratory Animal Co., Ltd. (Shanghai, China). Animals were housed in the specific pathogen-free (SPF-) grade animal facility at the School of Pharmacy, Soochow University, under controlled environmental conditions (temperature 22 ± 1 °C, relative humidity of 50 ± 10%, 12 - h light/dark cycle, and automatic ventilation) in compliance with the Guide for the Care and Use of Laboratory Animals. Mice were euthanised by cervical dislocation performed by trained personnel prior to tissue collection.

### 2.16 Mouse primary immune cell isolation

Bone marrow-derived dendritic cells (BMDCs) were isolated from 6-week-old C57BL/6 mice according to the protocol outlined by Sauter et al. (29). Cells were cultured in RPMI-1640 medium supplemented with 10% FBS, 1% P/S, 2 mM L-glutamine, 50 mM β-mercaptoethanol, and 20 ng/mL GM-CSF for 6-7 days. Splenocytes were isolated by mechanical dissociation and red blood cell lysis buffer (Solarbio, Cat# SLB-R1010, Beijing, China).

### 2.17 Immune cell functional assays

DC2.4, Raw264.7, and BMDCs or splenocytes (1.0 x 10^6^) were seed in 96-well plates and treated with purified sEVs (8 µg per well on day 0 and 2). On day 4, immune phenotypes were analysed by flow cytometry and cytokine secretion was quantified by Enzyme-linked immunosorbent assay (ELISA).

### 2.18 Flow Cytometry

To distinguish live cells, the Zombie Aqua™ Fixable Viability Kit (BioLegend, Cat#423102, San Diego, CA, USA) was used, and cells were incubated for 30 minutes. Next, Fc receptor was blocked for 5 min to minimise nonspecific antibody binding. Following that, the cells were incubated with fluorochrome-conjugated primary antibodies specific cell surface markers (Supplementary Table S3), according to the manufacturer’s recommended dilution, for 30 minutes in dark. Intracellular markers were detected after fixation/permeabilisation. Activated DCs were identified by expression of CD11c^+^, CD86^+^, and CD80^+^; M1 macrophages were characterised as CD11c^-^, F4/80^+^, and CD86^+^; M2 macrophages were identified as CD11c^-^, F4/80^+^, and CD206^+^; And T regulatory cells Tregs were characterised by CD45^+^, CD3^+^, CD4^+^, CD25^+^, and Foxp3^+^. Data were acquired on a Beckman Cytoplex, recording 10,000 events per sample, and analysed by FlowJo software V10.

The list of antibodies used in this study are provided in Table 1.

**Table 1:**
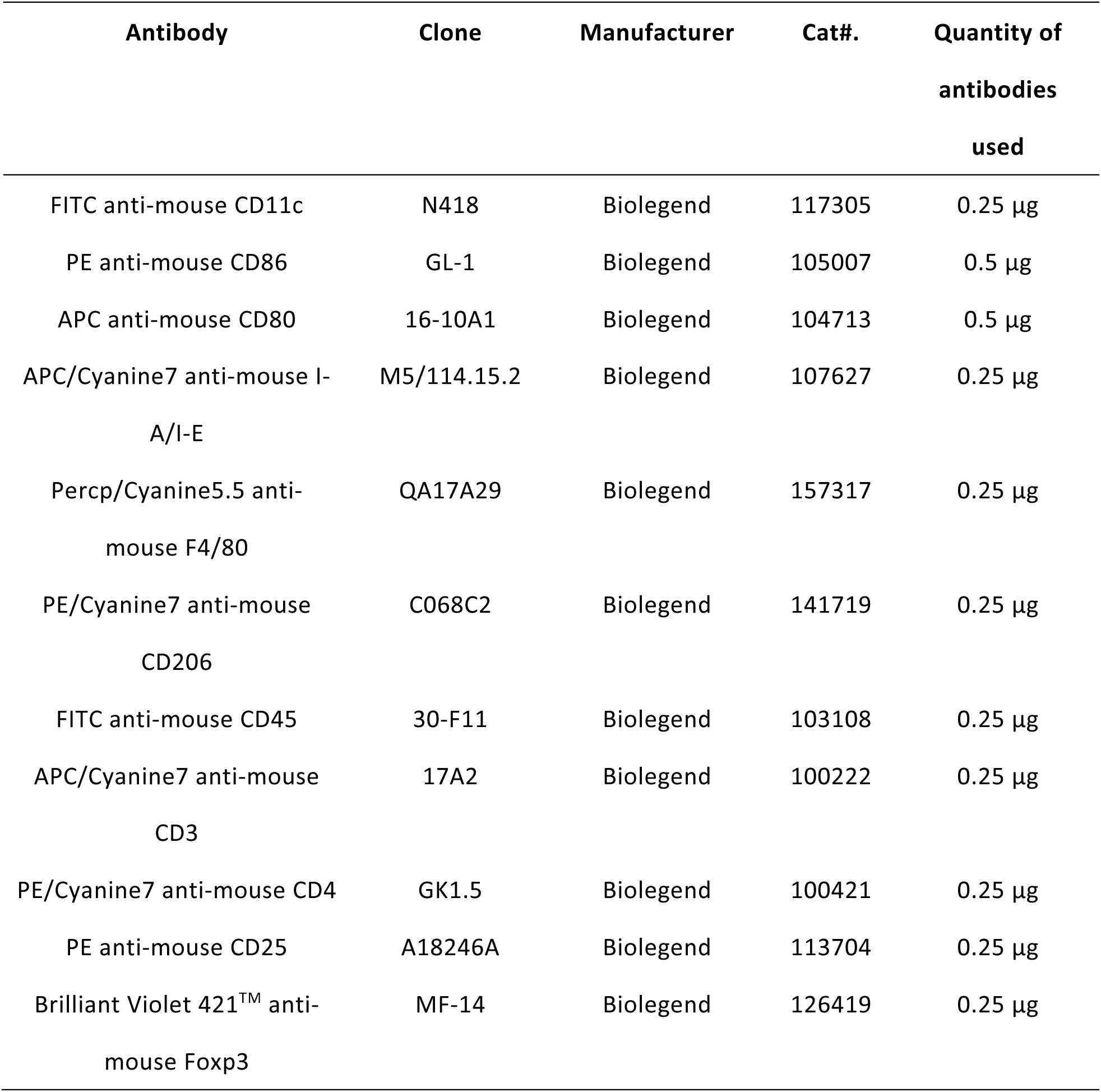
Fluorophore-conjugated antibodies for Flow Cytometry.

### 2.19 ELISA

Cytokine levels, including IL-6, IL-10, IL-12, TNF-α, and IFN- γ, secreted by BMDCs or Splenocytes following treatment with sEVs, were measured using ELISA. Commercial mouse ELISA kits (Invitrogen^TM^; IL-6, IL-10, IL-12p70, TNF alpha, IFN-γ) were used according to the manufacturer’s protocols. Briefly, an ELISA plate (Corning^TM^ Costar 9018, USA) was coated with capture antibody, blocked, and incubated with standards or samples, followed by detection antibody and Avidin-HRP. The reaction was developed with TMB substrate and stopped with acid solution. Absorbance was measured at 450 nm using a microplate reader.

### 2.20 Metabolites Analysis

Intracellular pyruvate and acetyl-CoA were measured using the Pyruvate Assay Kit (Solarbio, Cat#BC2205, Beijing, China) and Acetyl-CoA Content Assay kit (Geruisi, Cat#G0826W, Suzhou, China), respectively, in accordance with the manufacturer instructions. Lactate concentration in conditioned media was determined using a Lactic Acid Content Assay Kit (Solarbio, Cat#BC2235, Beijing, China).

### 2.21 Statistical analysis

Data are presented as the mean ± standard deviation (S.D.) and analysed using a student’s t-test to assess statistical significance. The analysis was performed with GraphPad Prism (GraphPad Software, CA, USA, version 10.1.2). For two group comparisons, tests were unpaired and two-tailed, with a statistical significance set at a p-value of < 0.05. While ANOVA test was used when multiple groups were involved. The p-value significance levels are as follows; ns (not significant) = p value ≥ 0.05, * = p value ≤ 0.05, ** = p value ≤ 0.01, *** = p value ≤ 0.001 and **** = p value ≤ 0.0001. All experiments were replicated independently, and the number of replicates is indicated in the figure legends.

## 3. Results

### 3.1 SK-M derived sEVs are enriched with MYCN

After stable transfection and single-cell cloning, SK-M cells exhibited over a 3.6-fold increase in MYCN protein and a 34-fold increase in mRNA levels. *MYCN* overexpression significantly enhanced cell proliferation and viability, consistent with its oncogenic functions(30)

(Supplementary Figure 1). Next, we verified enrichment of MYCN in sEVs secreted by SK and SK-M cells. Using differential ultracentrifugation, sEVs were isolated from SK and SK-M cells followed by NTA, TEM and WB according to MISEV 2023 guidelines(16). TEM analysis revealed typical spherical particles with a bright outer ring and a darker central area, while NTA analysis confirmed size distribution of these sEV samples from both cell types, with mean diameters of 141.1±3.20 nm (SK sEV) and 137.2±5.02 nm (SK-M sEV) (Figure 1C). WB demonstrated enrichment of exosome-associated markers (CD9 and HSP70) and the absence of the cellular marker (VDAC-1) in sEV fractions. Notably, MYCN was selectively enriched in sEVs derived from MYCN-overexpressing SK-M cells, consistent with its elevated expression in the corresponding cellular lysate (Figure 1D-E and Supplementary Figure 2).

**Figure 1:**
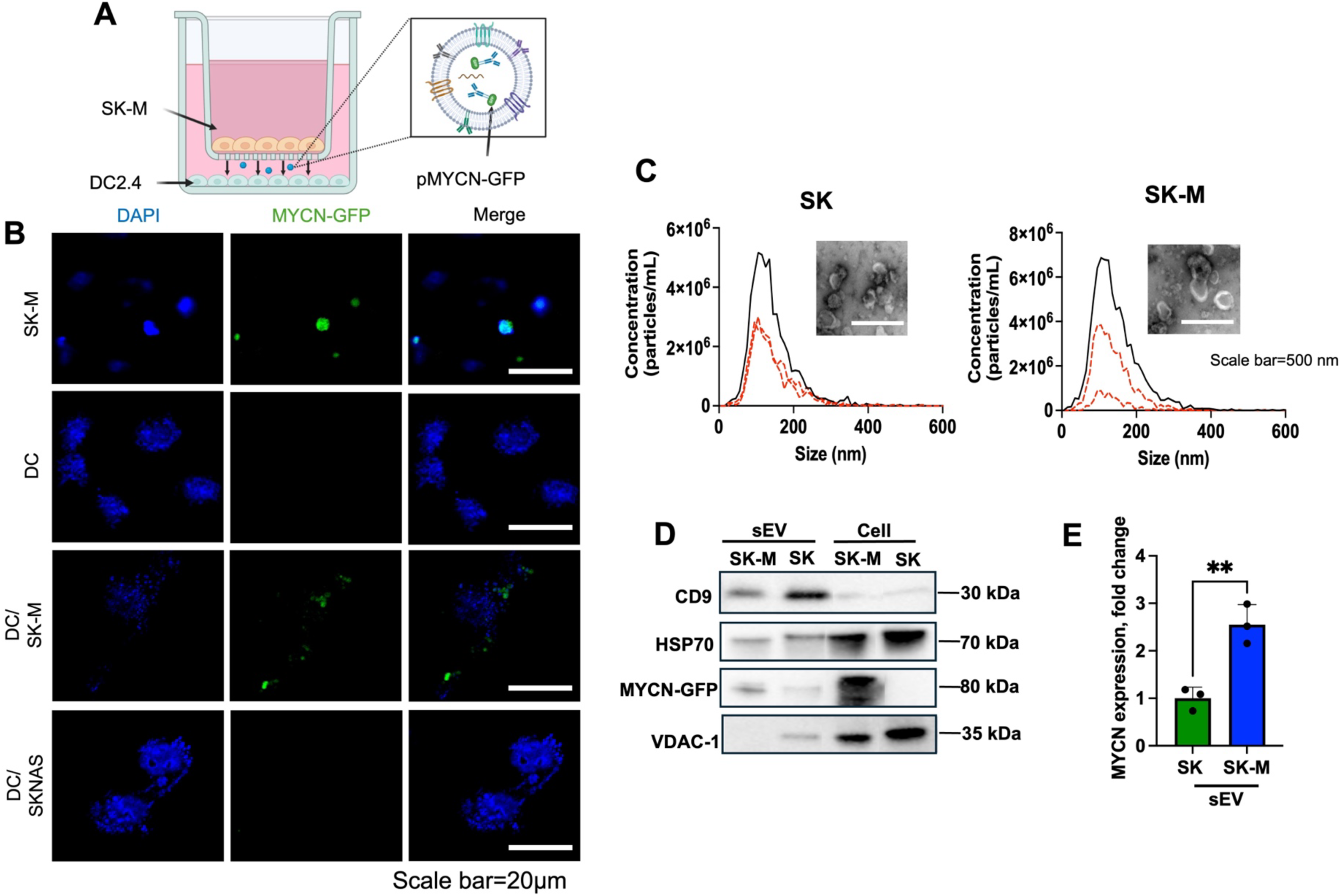
Isolation and Characterisation of sEV. (A) Diagram illustrating the transwell co-culture setup: SKNAS-*MYCN-GFP* (SK-M) cells were plated in the upper transwell insert, while DC2.4 cells were cultured in the lower compartment. (B) Representative fluorescence images of SK-M, DC2.4 and DC2.4 co-cultured with either SK-M or parental SKNAS cells, stained with Hoechst to label nuclei (N=3, N represents biological replicates of cells/sEVs). Scale bar = 20 µm. (C) Representative TME micrographs (top) and NTA size distribution profiles (bottom) of sEV isolated from SK and SK-M (N=3). Scale bar=500 nm. (D) Representative Western blot, of three independent experiments, showing the expression of CD9, HSP70, MYCN, and VDAC-1 in SK and SK-M cells and their sEVs, with densitometric quantification of MYCN levels in sEVs (E). Data represent mean ± standard deviation (SD) from three biological replicates. Statistical analysis was performed using an unpaired t-test (** = p value≤ 0.01).

To examine MYCN transfer, we set up a transwell co-culture system (0.4 µm pore size), which permits only soluble factors and sEVs to exchange between chambers. SK-M or SKNAS cells were seeded in the upper insert, and DC2.4 cells in the lower chamber. GFP signals detected in DC2.4 cells indicated potential MYCN-GFP transfer via sEVs (Figure 1A-B).

### 3.2 Functional enrichment analysis of sEV proteome in SK and SK-M

To further characterise the protein composition of SK and SK-M sEVs and evaluate proteomic alterations associated with *MYCN* amplification, we performed LC-MS/MS analysis followed by bioinformatic processing using Enrichr. Data were filtered, imputed, and subjected to differential expression analysis using SK cells/sEVs proteome as baseline.

Proteins with a fold change of >=1.5 and p-value <0.05 were considered upregulated, while those with a fold change of <=-1.5 and p-value <0.05 were defined as downregulated. Based on these criteria, 221 proteins were upregulated and 360 downregulated in SK-M cells (Supplementary Figure 3A). The 10 most enriched genes in SK-M are listed in Supplementary Table S4, and pathway enrichment analysis revealed a significant association with *MYC/MYCN* target gene networks (Supplementary Figure 3B-C).

Cross-referencing the cellular and sEV proteomes revealed that 83% of proteins in SK sEVs and 93% in SK-M sEVs overlapped with their respective parental cells, supporting the notion that sEVs largely reflect the protein repertoire of their cells of origin (Figure 2A).

**Figure 2:**
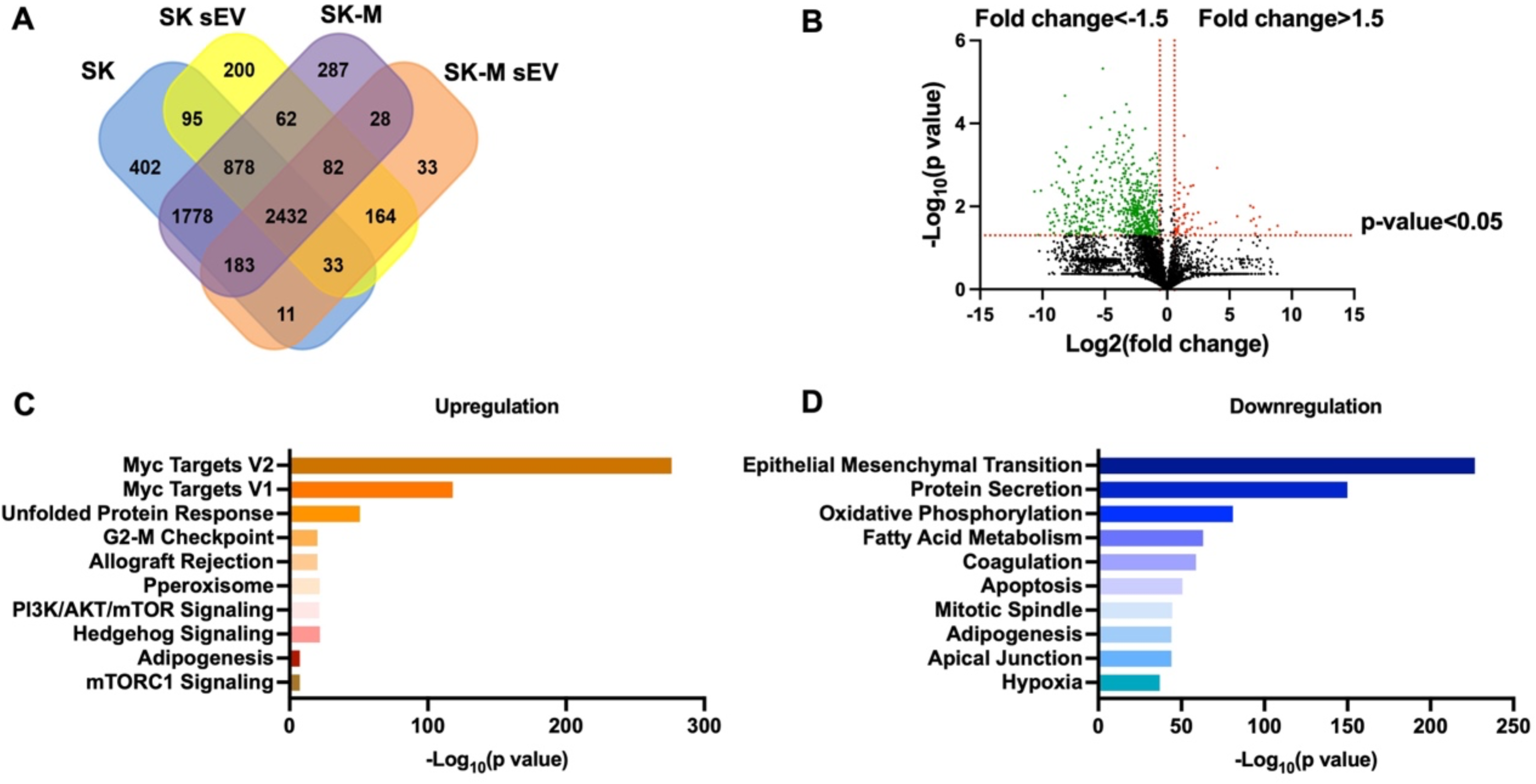
Mass spectrometry analysis of SK and SK-M sEV. (A) Venn diagram showing the overlap of total proteins identified in SK and SK-M sEVs and their corresponding parental cells. (B) Volcano plot displaying differential protein expression between SK and SK-M sEVs. Proteins shown in red are upregulated (fold change >1.5, p-value < 0.05), and those in blue are downregulated (fold change <-1.5, p-value < 0.05) in SK-M sEVs. Gene ontology (GO) enrichment analysis of significantly upregulated (C) and downregulated (D) proteins in SK-M sEVs relative to SK, ranked by –Log10(p-value) using Enrichr. LC-MS/MS analysis was performed on three biological replicates.

However, comparison of the top 10 upregulated proteins in SK-M cells and their corresponding sEVs identified distinct sets of 20 proteins, suggesting that sEVs selectively incorporate specific oncogenic cargos rather than passively mirroring the cellular proteome (Supplementary Table S4).

Functional enrichment analysis highlighted significant downregulation of OXPHOS and fatty acid metabolism pathways in SK-M sEVs, accompanied by upregulation of mTORC1 signaling (Figure 2B-D). These findings indicate that *MYCN* amplification induces profound metabolic reprogramming in sEVs, consistent with *MYCN*-driven metabolic traits such as enhanced glycolysis, lipid turnover, and nucleotide biosynthesis to support tumour cell proliferation(14).

### 3.3 sEVs secreted by SK-M cells increased lactate secretion immune cells

We further examined the glycolytic pathway as a representative marker of metabolic dysregulation (Figure 3A-F, Supplementary Figure 4 and Supplementary Figure 5). MYCN overexpression in SK-M cells and their sEVs led to enrichment of glycolytic enzymes compared with the non-*MYCN* amplified counterpart, in agreement with the previous studies by Tsakaneli et.al.(15) and Frawley et.al(31).

**Figure 3:**
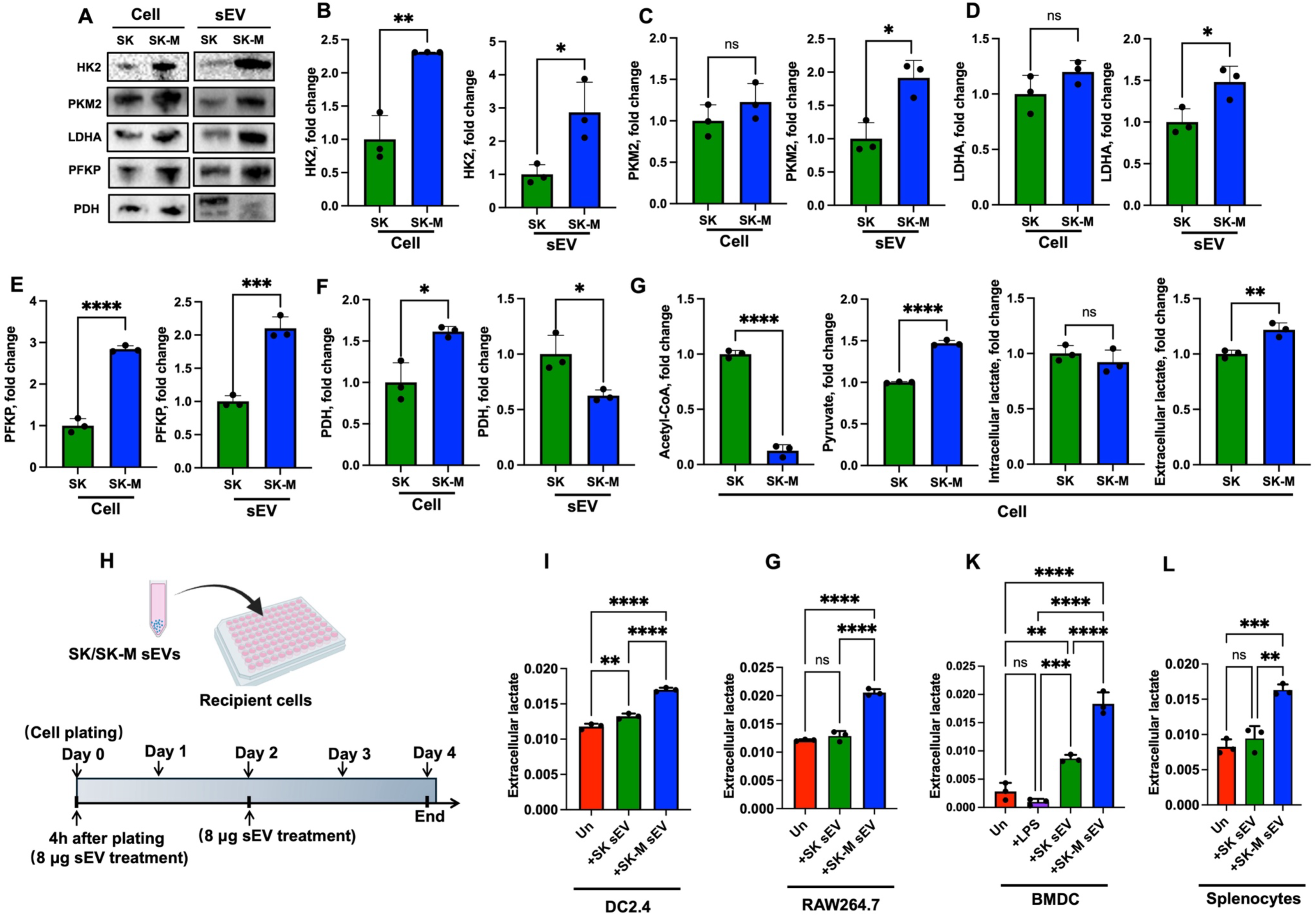
***MYCN* expression enhances glycolytic enzyme levels in SK-M cells and sEVs.** (A) Representative Western blot (from three independent experiments) showing the expression of HK2, PKM2, LDHA, and PFKP in SK and SK-M cells and their corresponding sEVs. Images are derived from separate blots prior to primary antibody incubation. (B-F) Densitometry analysis of HK2 (B), PKM2 (C), LDHA (D), PFKP (E), and Pyruvate Dehydrogenase (PDH) (F) in cellular lysates and sEV, represented as fold change relative to SK cells or sEVs after normalisation to endogenous controls or total protein staining, respectively. (G) Metabolic profiling of SK and SK-M cells showing key metabolic alterations, including elevated lactate secretion, represented as fold change relative to SK cells after normalisation to total protein content. (H) Schematic overview of the sEV treatment setup. On Day 0, immune cells were plated in 96-well plates and treated with the first dose of sEVs, followed by a second treatment on Day 2. Functional assays were performed on day 4. (I-L) Lactate secretion measured 96 hours after treatment with 16 µg of SK or SK-M sEVs in DC2.4 (I), Raw264.7(J), BMDC(K), and splenocyte (L) cultures, normalised to total protein content. “Un” denotes untreated control. Data are presented as mean ± SD from three biological replicates. Statistical significance was determined using an unpaired t-test (ns= not significant, ** = p value≤ 0.01, *** = p value≤ 0.001, and ****=p value≤ 0.0001

To further assess these metabolic alterations, we analysed the metabolite profiles of SK and SK-M cells. When cultured under identical conditions for 96 hours, SK-M cells exhibited a 21.8% increase in lactate secretion and an 87.4% reduction in acetyl-CoA levels compared with SK cells, indicating a metabolic shift toward glycolysis (Figure 3G).

Next, we examined whether sEV’s cargo could modulate lactate secretion in recipient cells within the immune NB microenvironment. Murine immune cell lines (DC2.4 and Raw264.7) and primary immune cells (BMDC and splenocyte) were treated with SK or SK-M sEVs (Figure 3H). After 96 hours, SK-M sEV treatment increased lactate secretion by 35.58%, 66.15%, 112.3%, and 72.87% in DC2.4, Raw264.7, BMDC, and splenocyte, respectively, compared with SK sEV treatment (Figure 3I-L). These findings suggest that SK-M sEVs can reprogram immune cell metabolism toward a more glycolytic phenotype by promoting lactate production and secretion, thereby increasing extracellular lactate levels and altering the pH of the TME.

### 3.4 SK-M sEV Reprogram BMDC and Splenocyte Responses Toward Immune Tolerance

Given that disordered metabolic states characterised by hypoxia and elevated metabolites, particularly lactate, contribute to immunosuppression within the TME(25), we next evaluated whether SK-M-derived sEV modulate the phenotypes of BMDCs and splenocytes following 96 hr treatment. Flow cytometry revealed that SK-M sEV-treated BMDCs exhibited a significant upregulation of both activation markers compared to untreated cells, indicating activation. However, the extent of activation was lower than that triggered by LPS, indicating a moderate immunostimulatory effect. No significant difference was observed between the SK and SK-M sEV-treated groups (Figure 4A and Supplementary Figure 6). Cytokine profiling by ELISA indicated that SK-M sEVs increased IL-12 and IL-6 levels relative to untreated group but, compared with SK sEVs, induced a 42.46% reduction in IL-12 production and a 316.9% increase in IL-10 secretion (Figure 4B). This cytokine pattern-elevated IL-10 concurrent with reduced IL-12-is indicative of an immunoregulatory or tolerogenic DC phenotype that supports immune tolerance, favors regulatory T cell differentiation, and suppresses cytotoxic T cell responses, despite evidence of partial activation(32, 33). The increased proliferation of BMDCs following sEV treatment further supports an immature or semi-mature phenotype, as mature DCs typically exit the cell cycle (Supplementary Figure 7A-B).

**Figure 4:**
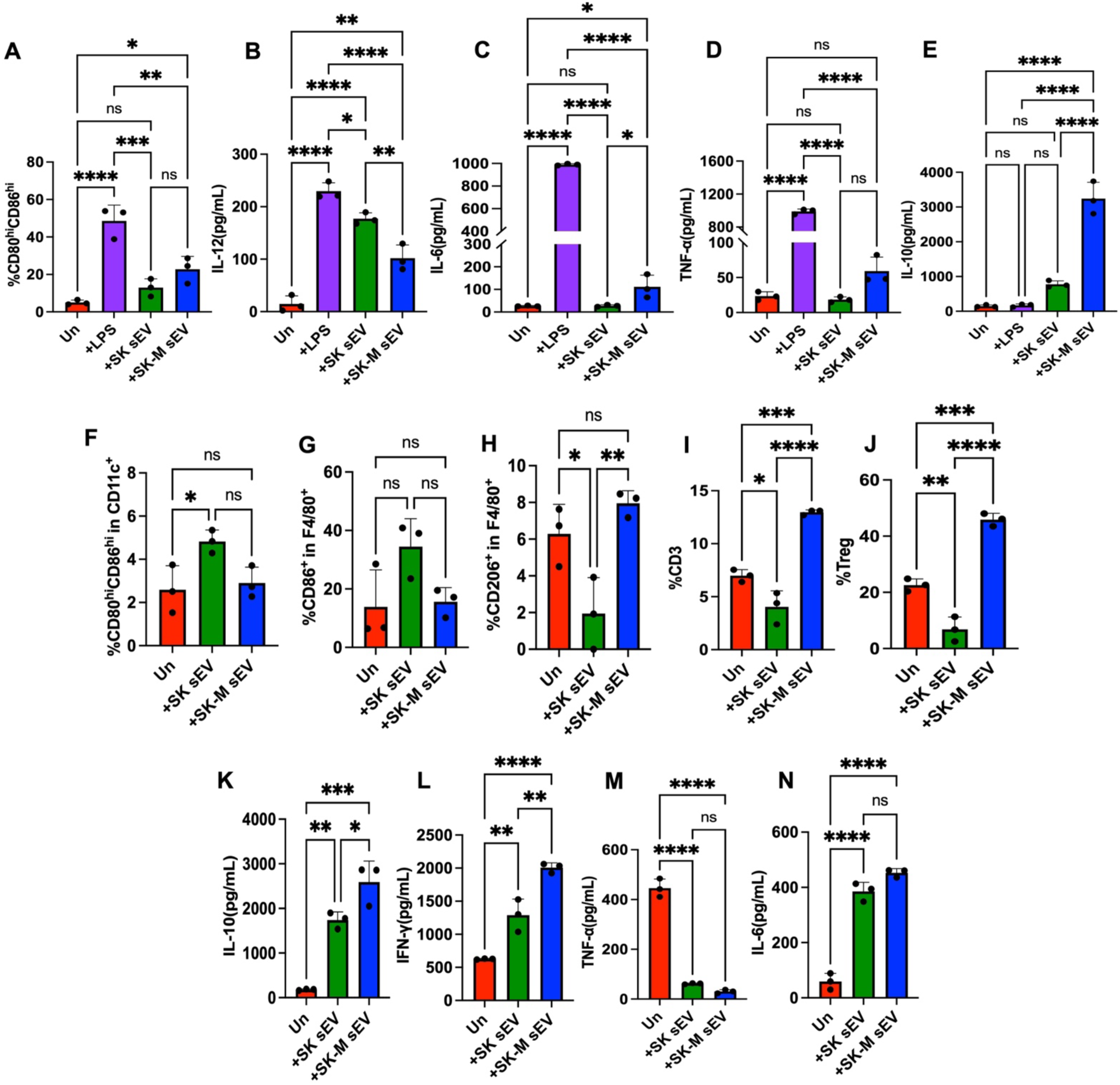
SK-M sEVs promote immunosuppression in BMDCs and splenocytes. (A) Activation of BMDCs assessed 4 days post-treatment with 16 µg of SK or SK-M sEVs, based on CD80 and CD86 expression. (B-E) Cytokine secretion by BMDC measured by ELISA 4 days after treatment with 16µg of SK or SK-M sEVs. (F-J) Flow cytometry analysis of splenocytes 4 days post-treatment with 16µg of SK or SK-M sEVs: DC activation based on CD80 and CD86 expression; (G-H) macrophage polarisation evaluated by CD86 (M1 marker) and CD206 (M2 marker) expression; (I-J) quantification of T cells particularly Tregs. (K-N) Cytokine secretion by splenocytes measured by ELISA 4 days post-treatment with 16µg of SK or SK-M sEVs. Graphs represent mean ± SD from three biological replicates. Statistical significance was calculated using an ordinary one-way ANOVA (ns = p > 0.05, * = p ≤ 0.05, ** = p ≤ 0.01, *** = p ≤ 0.001, **** = p ≤ 0.0001).

In splenocytes, a heterogeneous population comprising T lymphocytes, B cells, natural killer (NK) cells, macrophages, DCs, SK-M sEVs treatment significantly increased the total T cell population, particularly Tregs, and promoted macrophage polarisation from a pro-inflammatory M1 to a more “tumour-friendly” M2 phenotype (Figure 4C-E and Supplementary Figure 6). Consistently, SK-M sEVs induced a 49.22% increase in IL-10 secretion compared with SK sEV (Figure 4F). In proliferation assays, splenocytes treated with sEV exhibited reduced proliferative capacity relative to untreated and LPS-positive controls; however, no statistically significant difference was observed between the two sEV-treated groups (Supplementary Figure 7C-D).

## 4. Discussions

MYCN is the most well-characterised oncogenic driver and a key indicator of poor prognosis in neuroblastoma, playing an essential role in clinical risk stratification(8, 34). Beyond its canonical functions in proliferation and differentiation blockade(10, 35, 36), MYCN profoundly reshapes tumour cell physiology by regulating cellular stress responses, metabolic reprogramming, and adaptive survival pathways(9, 14, 15, 37).

For example, MYCN activation induces oxidative and proteotoxic stress, necessitating compensatory mechanisms such as the unfolded protein response (UPR) to sustain tumour cell viability(37). Additionally, it drives a metabolic shift towards aerobic glycolysis, glutamine dependence, and altered mitochondrial dynamics, supporting rapid biomass accumulation and redox homeostasis(9, 11, 14, 15)

Recent studies have also documented the ability of sEVs to reprogramme the cells within the neuroblastoma microenvironment by delivering oncogenic content such as miRNA(38, 39), MYCN protein itself(15). Our recent study reported that the MYCN oncogene was enriched in sEVs produced by NB cells with MYCN amplification(31).

Tsakaneli et.al. proposed a novel mechanism of non-cell-autonomous metabolic regulation, whereby *MYCN*-overexpressing NB cells “broadcast” metabolic reprogramming signals to neighbouring or distant cells through sEVs enriched in glycolytic enzymes, thereby enhancing glycolysis, and promoting the NB progression and metastasis(15). However, their study neither accounts for potential off-target effects of doxycycline on cellular metabolism at commonly used concentrations (100 ng/mL to 5 µg/mL) inherent to the SHPE-Tet/21N system nor investigates the contribution of MYCN-enriched EVs into noncancerous cells of the neuroblastoma microenvironment, including immune cells (40).

In this study, we utilised a different in vitro model of constitutively *MYCN*-expressing NB (SKNAS-MYCN-GFP, SK-M) developed in our lab to visualise *MYCN* localisation and intercellular transfer, characterise the sEV proteome, assess associated phenotypic changes, and investigate the potential role of sEVs in mediating *MYCN* transfer and metabolic reprogramming in the recipient non-cancerous immune cells. We confirmed enrichment of sEVs with MYCN and demonstrated that sEVs transfer MYCN to the recipient cell nucleus in a co-culture system, suggesting that they can regulate gene expression as a ‘guest’ transcription factor and remodel cellular behaviour within the TME.

The large number of differentially expressed proteins in SK-M sEV indicates a transformative potential for the recipient cell machinery once sEV content is released. A comparative analysis of the cellular and sEV proteomes revealed that sEVs selectively incorporate oncogenic cargos, rather than passively reflecting the cellular proteome. In SK-M sEVs, most downregulated pathways were associated with metabolism, including OXPHOS (Supplementary Table S5), indicating metabolic dysregulation. Further validation of the proteomics and pathway analyses demonstrated that MYCN-overexpressing SK-M cells and their derived sEVs were enriched with key glycolytic enzymes; the same pattern was reported by Tsakaneli et al. and LDHA, PKM1/2 and PKM2 by Frawley et al(31). Consistently,

SK-M cells exhibited a >21.8% increase in lactate secretion compared with their parental non-*MYCN*-amplified counterparts, reflecting a metabolic shift toward glycolysis. Increasing evidence indicates that tumour cells undergo metabolic reprogramming to support survival and confer resistance to chemotherapy(41, 42). The metabolic plasticity, driven by oncogenes such as MYCN and influenced by the TME, enables tumour cells to switch between aerobic glycolysis and OXPHOS(15, 41).

sEVs have attracted increasing attention in NB as key mediators of communication in TME (20). Growing evidence suggests that sEV secreted by NB cells actively influence neighbouring stromal components, thereby facilitating tumour progression. Nakata et al. reported that NB-derived sEVs reprogram mesenchymal stem cells (MSCs), markedly enhancing their secretion of cytokines, such as VEGF, IL-17RA, IFN-α, IL-5, IL-6, IL-8, MCP-1, and MIP-1β(21). This cytokine shift promoted a tumour supportive MCS phenotype that contributed to NB dissemination(21). Similarly, Colletti et al. found that sEVs from metastatic NB cells were internalised by MSCs, leading to upregulation of osteogenic markers including BMP2, RUNX2, SPP1 and OSTERIX via miRNA miR-375. which enhanced the pro-metastatic behaviour of MSCs and increased tumour invasiveness(22).

These EVs contribute to immune suppression by enabling tumour immune evade, inducing immune cell apoptosis, and promoting resistance to immunotherapies, thereby transforming “hot” (immune-active) tumours into “cold” (immune-silent) ones (20, 24, 43). They can deliver oncogenic signals, remodel the TME, and modulate immune responses(20, 28, 44–52).

However, few studies have examined how *MYCN*-overexpressing NB cells reprogram immune cells within the TME through sEV-mediated signaling to promote immune escape. The metabolic and phenotypic characterisation of BMDCs and splenocytes in this study addressed this critical gap. Our findings demonstrate that sEVs derived from both SK and SK-M cells exert significant immunomodulatory effects, with distinct differences reflecting the influence of the *MYCN* oncogene. Specifically, SK-M sEVs enhanced lactate production and secretion in immune cells, promoting a more tumour supportive phenotype through the induction of immunosuppressive populations. The resulting elevation of extracellular lactate levels further acidified the TME, thereby fostering the expansion of inmmunosuppressive cell subsets such as M2 macrophages and Tregs.

### Study limitations

The potential role of sEVs in transferring the *MYCN* oncogene to immune cells warrants further investigation using multiple human immune cell models exposed to sEVs. Expanding this concept to human non-cancerous cells within TME would further elucidate the mechanisms of sEV-mediated signalling and their contribution to tumour progression and prognosis. Although SK-M sEV treatment significantly increased immunosuppressive BMDC and splenocyte populations, the biological effect size was modest. These observations should therefore be interpreted with caution, as the findings have not yet been verified in vivo models and/or clinical samples.

## Availability of data and materials

The mass spectrometry proteomics data have been deposited to the ProteomeXchange Consortium (https://www.iprox.cn/page/project.html?id=IPX0014697000) via the iProX partner repository with the dataset identifier PXD071984 (and iProX accession IPX0014697000). All data generated or analysed during this study are included in this published article and its supplementary information files.

## CRediT authorship contribution statement

L.M.: Conceptualisation; Formal analysis; Investigation; Methodology; Validation; Visualisation; Writing—original draft; Writing—review & editing.

M.L.: Resources, Project administration, Supervision, Funding Acquisition; Writing – review & editing.

O.P.: Conceptualisation; Data curation; Formal analysis; Funding acquisition; Investigation; Methodology; Resources, Project administration; Supervision; Visualisation; Writing—original draft; Writing—review & editing

## Supplementary information

Supplementary information to this article can be found online at

