## Supplemental File for "Small extracellular vesicular transfer of MYCN and glycolytic cargo coordinates metabolic and immunological reprogramming in neuroblastoma in vitro"

### Supplementary Information

**Supplementary Table S1*:* qPCR Primer Sequences.**

| Primer Name | Sequence (5’ to 3’) | Length | Cat no. | Manufacturer |
|---|---|---|---|---|
| Human MYCN Forward Primer | ACCCGGACGAAGATGACTTCT | 21 | S3121117465D | GENEWIZ |
| Human MYCN Reverse Primer | CAGCTCGTTCTCAAGCAGCAT | 21 | S3121117466D | GENEWIZ |
| Human GAPDH Forward Primer | GGAGCGAGATCCCTCCAAAAT | 21 | S3121117467D | GENEWIZ |
| Human GAPDH Reverse Primer | GGCTGTTGTCATACTTCTCATGG | 23 | S3121117468D | GENEWIZ |

**Supplementary Table S2*:* Primary antibodies for western blotting.**

| | Antibody | Species | Mw (kDa) | Manufacturer | Cat no. |
|---|---|---|---|---|---|
| EV characterisation | HSP70 | Rb | 72,73 | Cell Signalling Technology | 4872T |
| | VDAC-1 | Rb | 32 | Cell Signalling Technology | 4661S |
| | CD9 | Rb | 23-30 | Proteintech | 20597-1-AP |
| | MYCN | Rb | 62 | Cell Signalling Technology | 84406s |
| Endogenous control | β-Tublin | Rb | 55 | Cell Signalling Technology | 2128 |
| | GAPDH | Ms | 36 | Bio-Rad | MCA4740 |
| | β-actin | Rb | 45 | Cell Signalling Technology | 8457T |
| Glycolytic pathway | Hexokinase II | Rb | 102 | Cell Signalling Technology | 2867 |
| | PKM1/2 | Rb | 60 | Cell Signalling Technology | 3190 |
| | PFKP | Rb | 80 | Cell Signalling Technology | 8164 |
| | Pyruvate Dehydrogenase | Rb | 43 | Cell Signalling Technology | 3205 |
| | Hexokinase I | Rb | 102 | Cell Signalling Technology | 2024 |
| | PKM2 | Rb | 60 | Cell Signalling Technology | 4053 |
| | LDHA | Rb | 37 | Cell Signalling Technology | 3582 |

**Supplementary Table S3: Fluorophore-conjugated antibodies for Flow Cytometry.**

| Antibody | Clone | Manufacturer | Cat no. | Quantity of antibodies used |
|---|---|---|---|---|
| FITC anti-mouse CD11c | N418 | Biolegend | 117305 | 0.25 μg |
| PE anti-mouse CD86 | GL-1 | Biolegend | 105007 | 0.5 μg |
| APC anti-mouse CD80 | 16-10A1 | Biolegend | 104713 | 0.5 μg |
| APC/Cyanine7 anti-mouse I-A/I-E | M5/114.15.2 | Biolegend | 107627 | 0.25 μg |
| Percp/Cyanine5.5 anti-mouse F4/80 | QA17A29 | Biolegend | 157317 | 0.25 μg |
| PE/Cyanine7 anti-mouse CD206 | C068C2 | Biolegend | 141719 | 0.25 μg |
| FITC anti-mouse CD45 | 30-F11 | Biolegend | 103108 | 0.25 μg |
| APC/Cyanine7 anti-mouse CD3 | 17A2 | Biolegend | 100222 | 0.25 μg |
| PE/Cyanine7 anti-mouse CD4 | GK1.5 | Biolegend | 100421 | 0.25 μg |
| PE anti-mouse CD25 | A18246A | Biolegend | 113704 | 0.25 μg |
| Brilliant Violet 421$^{TM}$ anti-mouse Foxp3 | MF-14 | Biolegend | 126419 | 0.25 μg |

**Supplementary Table S4: Top 10 significantly upregulated proteins in SK-M sEV in comparison with parental cells identified by label-free LC-MS/MS.**

| Gene names | Protein names | Protein IDs | Cell | | | sEV | | |
|---|---|---|---|---|---|---|---|---|
| | | | Fold change | Significance | Modulation | Fold change | Significance | Modulation |
| ITGA10 | Integrin alpha-10 | O75578 | 1099 | * | ↑ | Not significant | - | X |
| ABCA3 | Phospholipid-transporting ATPase | Q99758 | 375 | * | ↑ | Not significant | - | X |
| CAPG | Macrophage-capping protein | P40121 | 372 | * | ↑ | Not significant | - | X |
| IQGA2 | Ras GTPase-activating-like protein | Q13576 | 300 | * | ↑ | Not significant | - | X |
| MYCN | N-myc proto-oncogene protein | P04198 | 277 | * | ↑ | 174 | * | X |
| ITPR3 | Inositol 1,4,5-trisphosphate receptor type 3 | Q14573 | 273 | * | ↑ | Not significant | - | X |
| PHGDH | D-3-phosphoglycerate dehydrogenase | O43175 | 267 | * | ↑ | Not significant | - | X |

**Supplementary Table S4** (continued).

| Gene names | Protein names | Protein IDs | Cell | | | sEV | | |
|---|---|---|---|---|---|---|---|---|
| | | | Fold change | Significance | Modulation | Fold change | Significance | Modulation |
| PPTC7 | Protein phosphatase PTC7 homolog | Q8NI37 | 217 | * | ↑ | Not significant | - | X |
| PALLD | Palladin | Q8WX93 | 189 | * | ↑ | 112 | ns | X |
| UTRN | Utrophin | P46939 | 179 | * | ↑ | 10 | ns | X |
| BDKRB1 | B1 bradykinin receptor | P46663 | Not significant | - | X | 1327 | * | ↑ |
| SPOCK1 | Testican-1 | Q08629 | Not significant | - | X | 464 | * | ↑ |
| NOC4L | Nucleolar complex protein 4 | Q9BVI4 | Not significant | - | X | 295 | * | ↑ |
| DUS3L | tRNA-dihydrouridine (47) synthase [NAD(P)(+)]-like | Q96G46 | 3 | ns | X | 172 | * | ↑ |

**Supplementary Table S4** (continued).

| Gene names | Protein names | Protein IDs | Cell | | | sEV | | |
|---|---|---|---|---|---|---|---|---|
| | | | Fold change | Significance | Modulation | Fold change | Significance | Modulation |
| MARF1 | Meiosis regulator and mRNA stability factor 1 | Q9Y4F3 | 15 | ns | X | 133 | * | ↑ |
| PIK3C2A | Phosphatidylinositol 4-phosphate 3-kinase C2 domain-containing subunit alpha | O00443 | 4 | ns | X | 123 | * | ↑ |
| POLR3A | DNA-directed RNA polymerase III subunit RPC1 | O14802 | Not significant | - | X | 120 | * | ↑ |
| HABP2 | Hyaluronan-binding protein 2 | Q14520 | 6 | ns | X | 106 | * | ↑ |
| RPF1 | Ribosome production factor 1 | Q9H9Y2 | Not significant | - | X | 102 | ** | ↑ |

Annotation: Significance is denoted as: ns = $p > 0.05$, * = $p \leq 0.05$, ** = $p \leq 0.01$. The symbol "↑" indicates upregulation and "X" indicates no significant difference when compared to SK cells or sEVs.

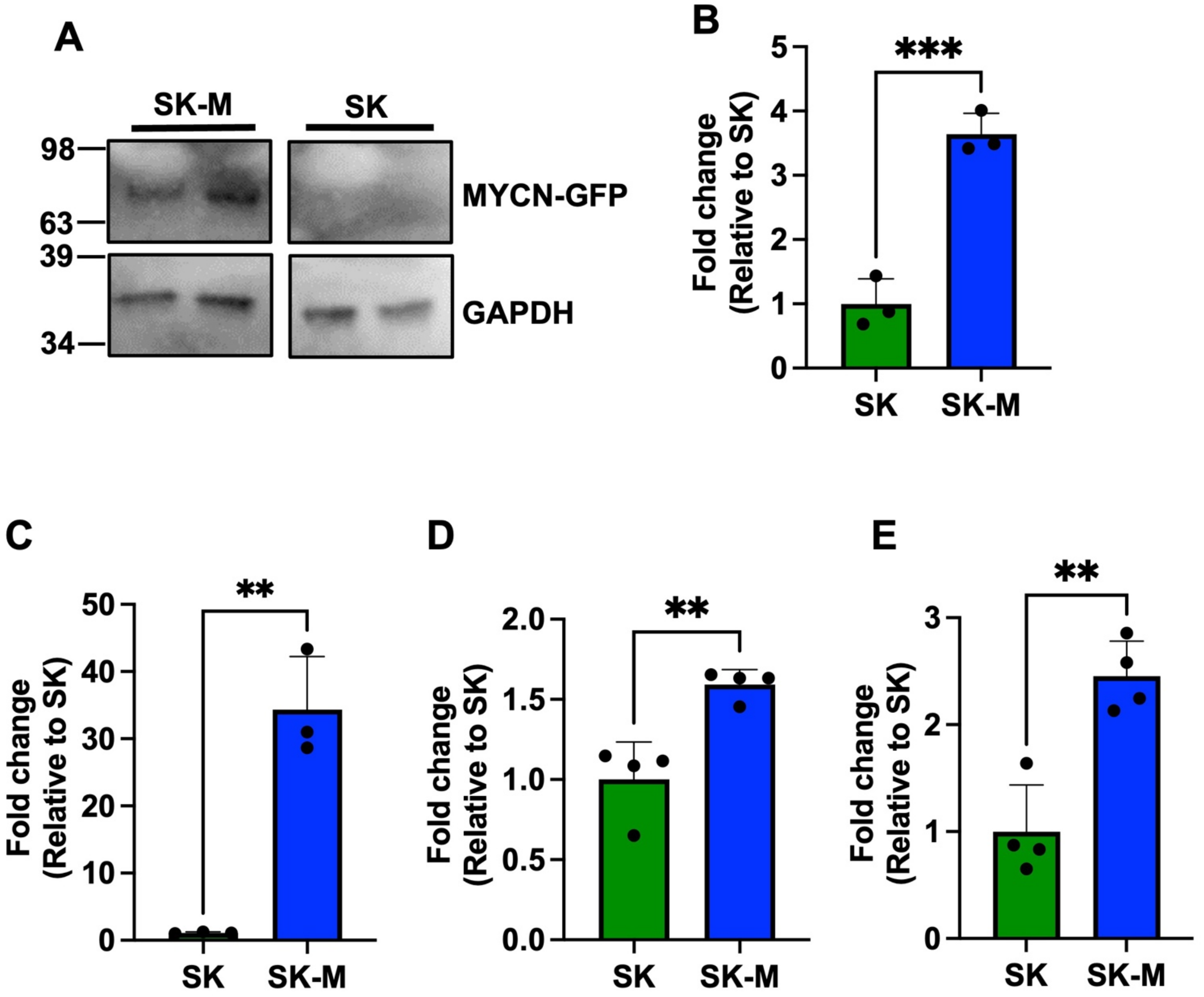


**Supplementary Figure 1: MYCN amplification in SKNAS cells induces cell proliferation and viability.** (A) Representative Western blot (from two independent experiments) showing MYCN expression in SKNAS-GFP (SK) and SKNAS-MYCN-GFP (SK-M) cells. (B) Densitometric quantification of MYCN expression in (A), presented as fold change relative to SK after normalisation to GAPDH. (C) Relative MYCN mRNA expression in SK-M compared with SK, normalised to GAPDH (N=3, N represents biological replicates of cells). (D) Cell proliferation and (E) cell viability of SK and SK-M cells, expressed as fold change relative to SK (N=4). Data are presented as mean ± SD. Statistical significance was calculated using an unpaired t-test (** = p value ≤ 0.01 and *** = p value ≤ 0.001).

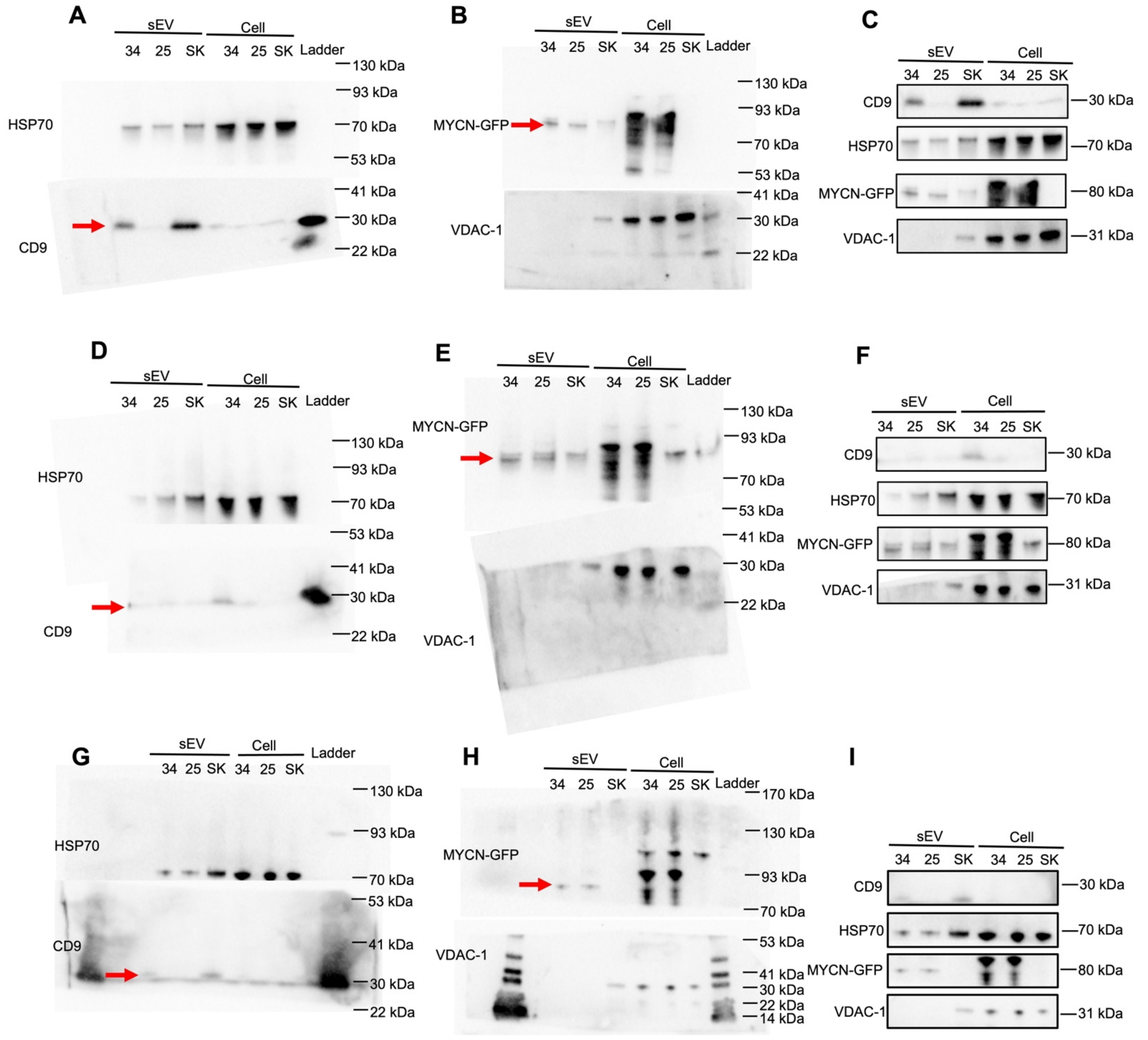


**Supplementary Figure 2: Uncropped full western blot WB images corresponding to Figure 1D.** The panel shows three independent biological replicates. Representative blots from these experiments are presented in Figure 1D. Blots are probed for exsomal markers CD9 and HSP70, as well as MYCN-GFP and mitochondrial marker VDAC-1 in SK and SK-M cells and their sEVs, Images for HSP70 and CD9 are derived from the same blot, cut into two, prior to primary antibody incubation. Similarly, MYCN-GFP and VDAC-1 were detected on another membrane that was also cut prior to primary antibody incubation. Clone 34 and 25 are two independent MYCN-transfected SKNAS clones; all experiments throughout the study were performed using clone 34 (designated SK-M).

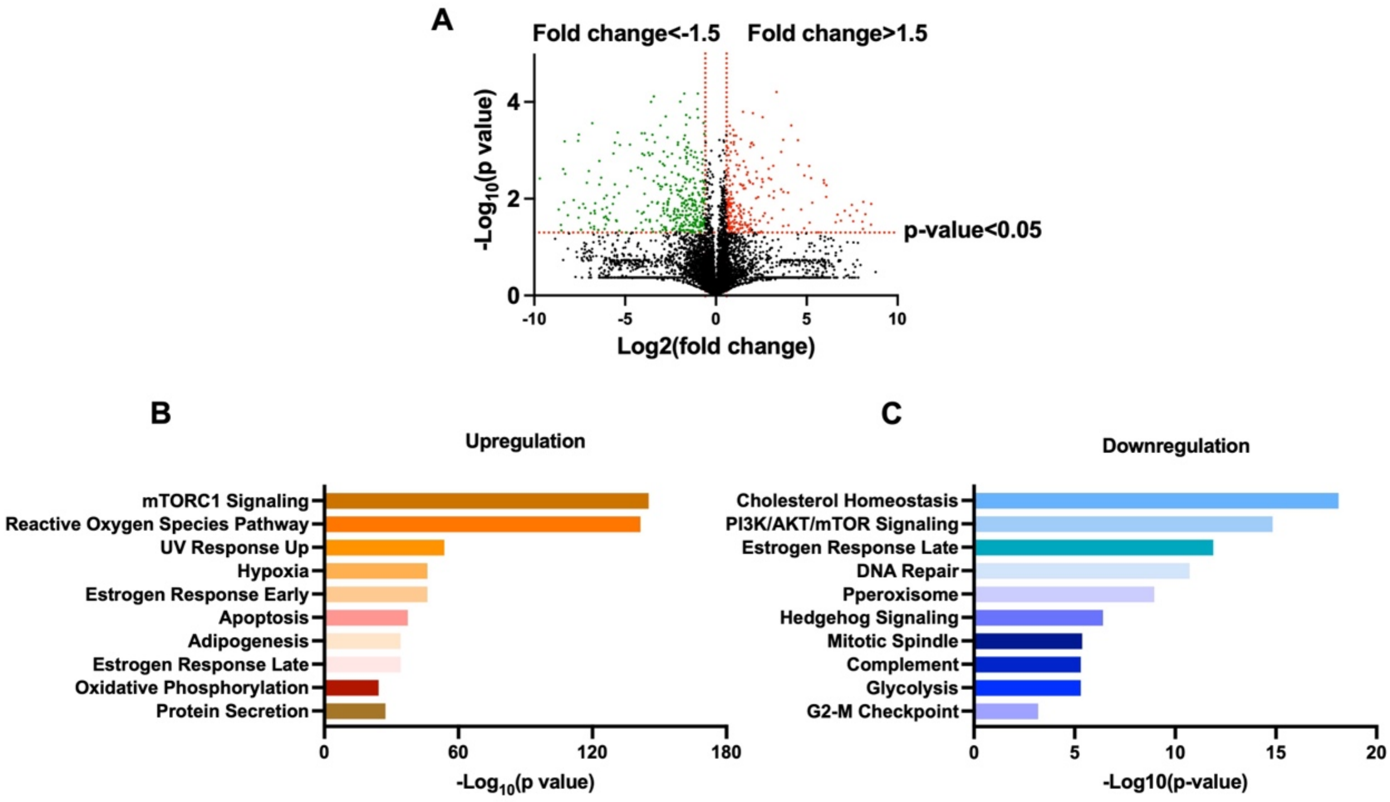


**Supplementary Figure 3: Proteomic profiles of SK and SK-M cells.** (A)Volcano plot depicting differentially expressed proteins between SK and SK-M cells. (B-C) Pathway enrichment analysis of proteins upregulated (B) and downregulated (C) in the SK-M cells, performed using Enrichr. All proteomic analyses were performed using three biological replicates.

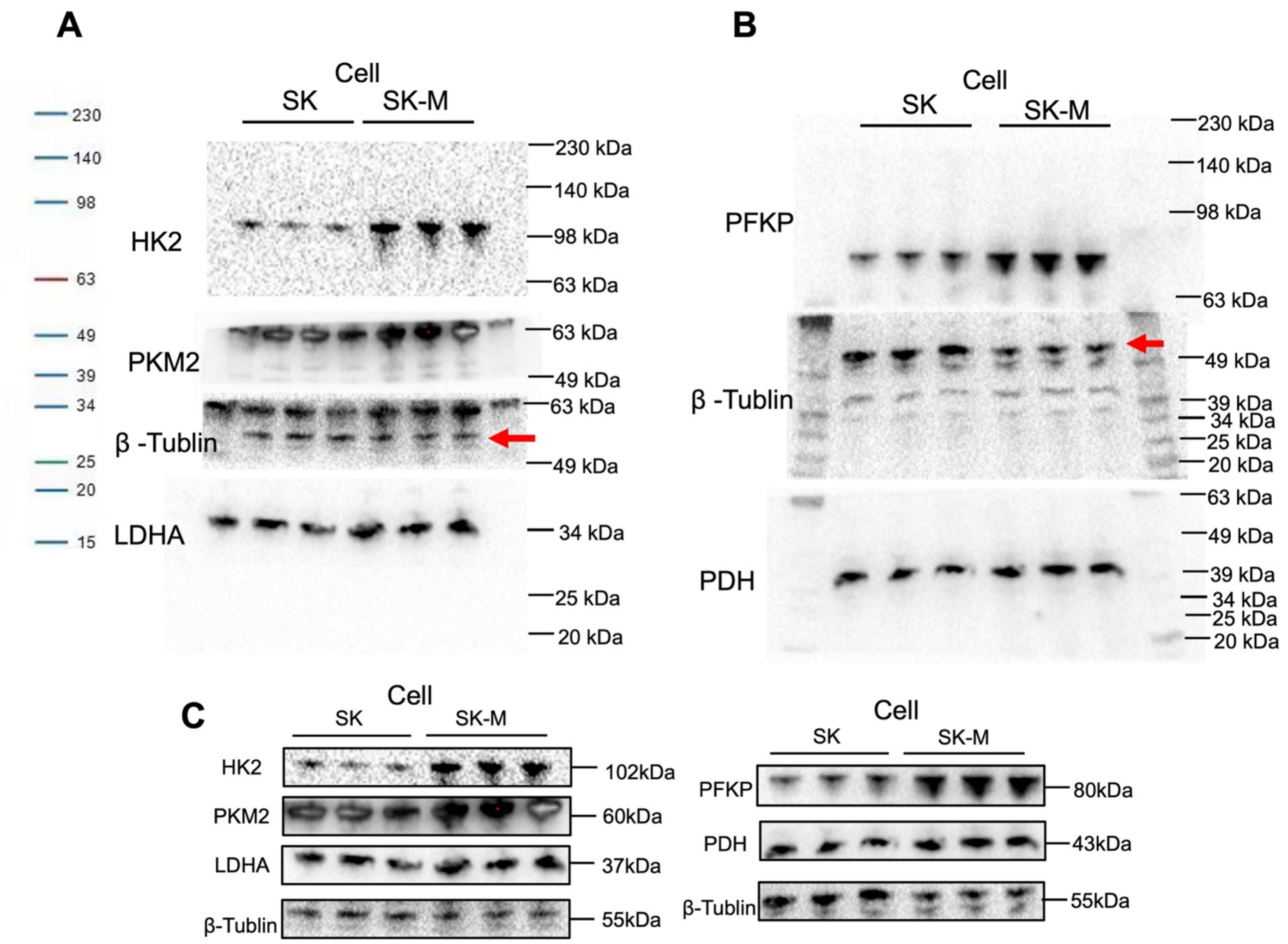


**Supplementary Figure 4: Uncropped full western blot WB images corresponding to Figure 3A.** The panel shows three independent biological replicates. (A-B) Blots are probed for glycolytic emzyme markers HK2, PKM2, LDHA, PFKP, Pyruvate Dehydrogenase (PDH) as well as β -Tublin in SK and SK-M cells. Images for HK2, PKM2, and LDHA are derived from the same blot, cut into three, prior to primary antibody incubation. Similarly, PFKP and Pyruvate Dehydrogenase (PDH) were detected on another membrane that was also cut prior to primary antibody incubation. β-Tublin were probed after PKM2 or PFKP using the same blots. Combined blots showed in (C).

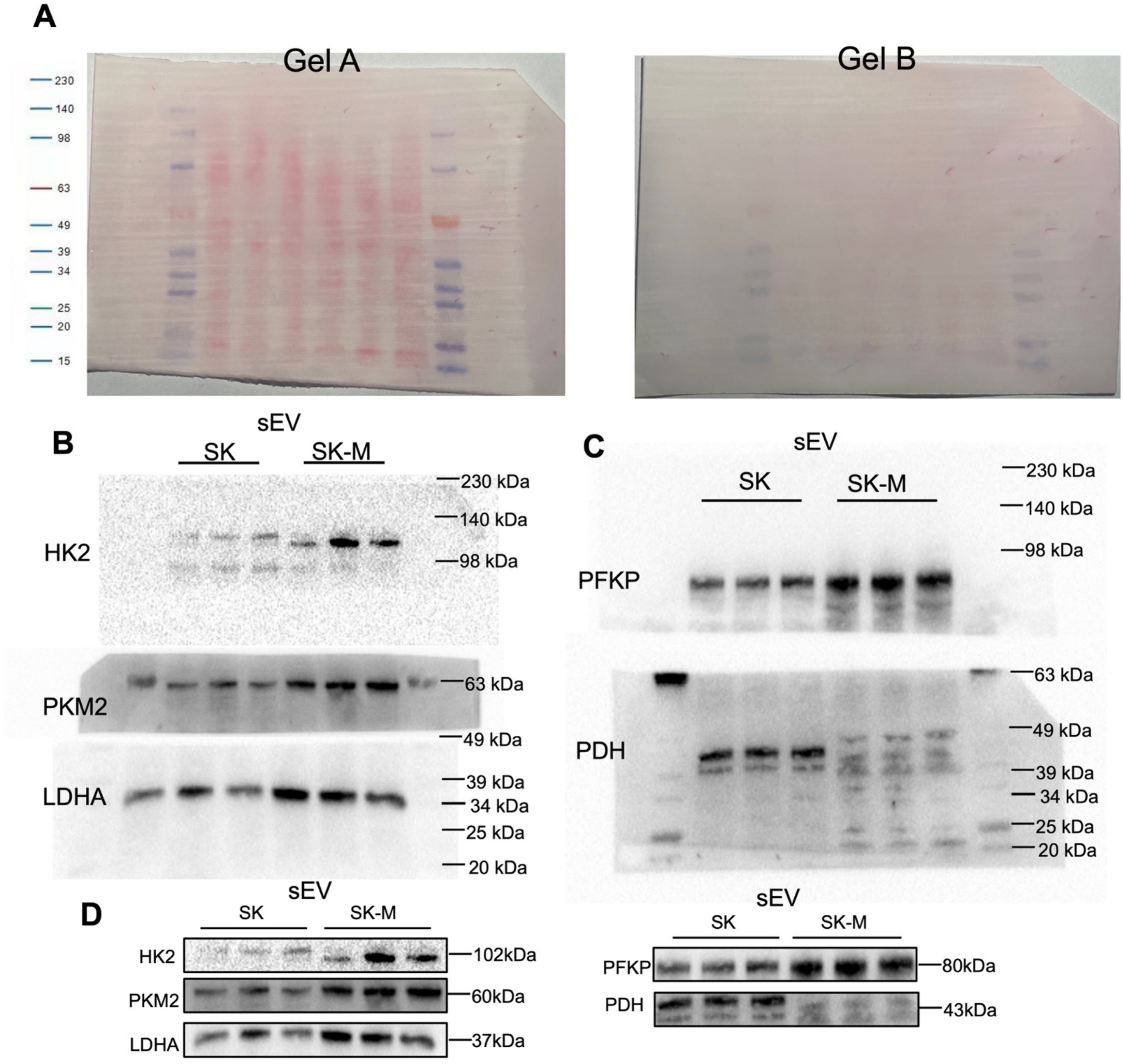


**Supplementary Figure 5: Uncropped full western blot WB images corresponding to Figure 3A.** (A) Stained membranes for immunodetection and normalization. (B-C) Blots are probed for glycolytic emzyme markers HK2, PKM2, LDHA, PFKP, Pyruvate, and PDH in SK and SK-M sEVs. Images for HK2, PKM2, and LDHA are derived from the same blot (Gel A), cut into three, prior to primary antibody incubation. Similarly, PFKP and PDH were detected on another membrane that was also cut prior to primary antibody incubation (Gel B). Combined blots showed in (D).

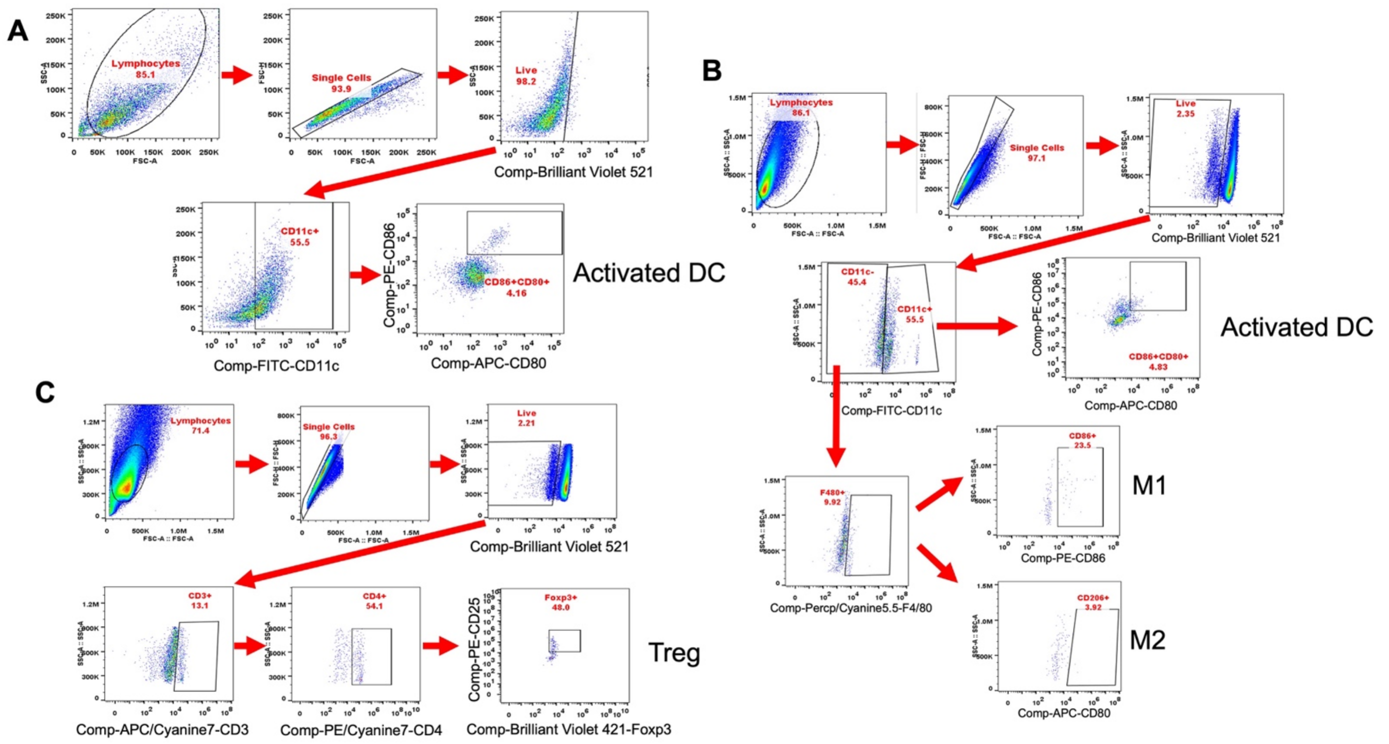


**Supplementary Figure 6: Gating strategies for flow cytometry analysis.** (A) Representative plots showing the gating strategy for activated dendritic cells (DCs) in BMDCs. (B-C) Representative plots showing the gating strategies for activated DCs, M1 (type 1) and M2 (type 2) macrophages (B), and Tregs (C) in the splenocytes. All flow cytometry analyses were performed in three biological replicates.

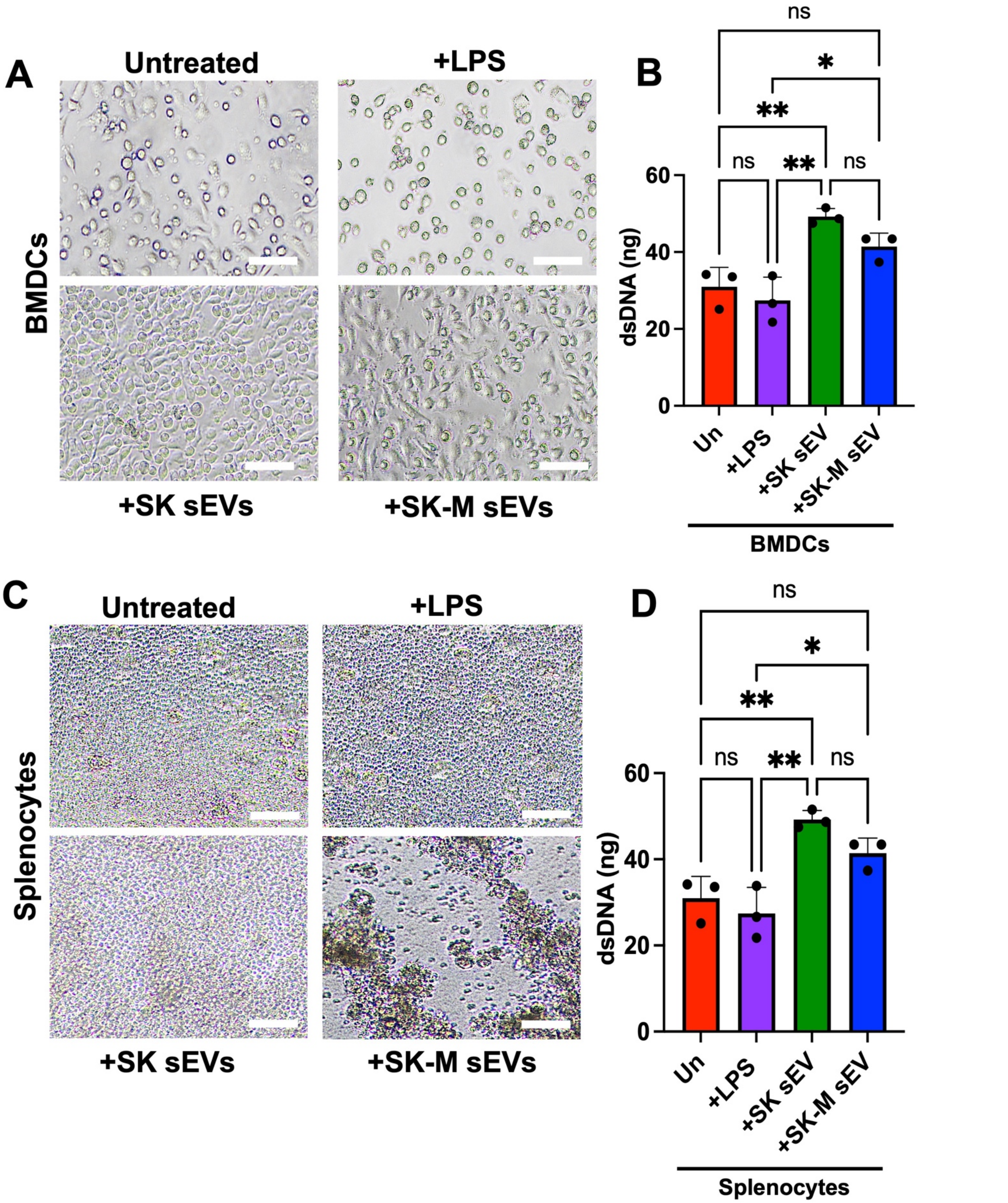


**Supplementary Figure 7: Cell proliferation of BMDCs and splenocytes 4 days post-treatment with 16 µg of SK or SK-M sEVs.** (A) Representative images of BMDCs after treatment (10x magnification; Scale bar=20 µm). (B) BMDCs proliferation quantified by dsDNA content 4 days post-treatment. (C) Representative images of splenocytes after treatment (10x magnification; Scale bar=20 µm). (D) Splenocytes proliferation quantified by dsDNA content 4 days post-treatment.

**Supplementary Table S5: Significantly upregulated proteins in SK-M sEV in comparison with SK sEV identified by label-free LC-MS/MS.**

| Gene names | Protein names | Protein IDs | Fold change | Significance | Modulation |
|---|---|---|---|---|---|
| NDUFA9 | NADH dehydrogenase [ubiquinone] 1 alpha subcomplex subunit 9 | Q16795 | 0.15943814 | * | ↓ |
| UQCRC1 | Cytochrome b-c1 complex subunit 1 | P31930 | 0.19452747 | * | ↓ |
| UQCRC2 | Cytochrome b-c1 complex subunit 2 | P22695 | 0.07660477 | ** | ↓ |
| IDH3A | Isocitrate dehydrogenase [NAD] subunit alpha | P50213 | 0.11256337 | * | ↓ |
| SUCLA2 | Succinate--CoA ligase [ADP-forming] subunit beta | Q9P2R7 | 0.0281972 | * | ↓ |
| CYB5A | Cytochrome b5 | P00167 | 0.00631701 | ** | ↓ |
| ATP6V1G1 | V-type proton ATPase subunit G 1 | O75348 | 0.01032415 | * | ↓ |
| SLC25A3 | Kidney mitochondrial carrier protein 1 | Q5SVS4 | 0.09991762 | * | ↓ |
| ACAA2 | 3-ketoacyl-CoA thiolase | P42765 | 0.04145968 | ** | ↓ |
| ATP6AP1 | V-type proton ATPase subunit S1 | Q15904 | 0.2390087 | * | ↓ |
| GPX4 | Phospholipid hydroperoxide glutathione peroxidase GPX4 | P36969 | 0.00712947 | ** | ↓ |
| MRPS22 | Small ribosomal subunit protein mS22 | P82650 | 0.01237415 | * | ↓ |
| IDH2 | Isocitrate dehydrogenase [NADP] | P48735 | 0.00678798 | * | ↓ |
| RHOT2 | Mitochondrial Rho GTPase 2 | Q8IXI1 | 0.02762265 | ** | ↓ |
| ATP5PO | ATP synthase subunit O | P48047 | 0.22213124 | * | ↓ |
| CYC1 | Cytochrome c1 | P08574 | 0.14783087 | ** | ↓ |
| DLD | Dihydrolipoyl dehydrogenase | P09622 | 0.17396547 | * | ↓ |
| ATP6V1C1 | V-type proton ATPase subunit C | P21283 | 0.16224687 | * | ↓ |
